# How to choose the optimal RNA-Seq library characteristics for alternative splicing analysis

**DOI:** 10.1101/2024.07.11.603071

**Authors:** Annika Ladwig, Melina Klostermann, Kathi Zarnack

## Abstract

Alternative splicing (AS) is a key layer of regulation in eukaryotic gene expression that is investigated in all areas of life sciences. Differences in AS between conditions can be quantified from transcriptome-wide short-read RNA sequencing (RNA-Seq) data with designated computational tools. However, not all short-read RNA-Seq data are equally suited for AS analysis. Here, we perform an exemplary AS analysis to showcase the impact of the RNA-Seq library characteristics on the obtained results. Using three standard ENCODE datasets with widespread AS changes, we modulate read length, read depth and the number of replicates and compare their influence on the detection, quantification and classification of AS events with the state-of-the-art AS algorithm MAJIQ. We find that longer reads and a higher read depth are the most effective measures to improve the sensitivity and precision of the analysis. From our results, we provide a recommendation on how to best choose the short-read RNA-Seq library specifications for an AS analysis.

## Introduction

Splicing is a universal process of RNA maturation in eukaryotes. During splicing, the noncoding introns are excised and the remaining exons are fused together to obtain a mature RNA. Intriguingly, most pre-RNAs can be spliced in multiple ways in a process called alternative splicing (AS), resulting in alternative RNA isoforms that utilize distinct exons or exon parts. Changes in AS play important roles in the regulation of gene expression, for instance during stem cell differentiation or cellular stress responses, and critically contribute to many human diseases including cancer (Jiang and Chen 2021). The detection of AS changes in various conditions has therefore become a topic of wide interest.

Different types of AS events can be discriminated based on the alternative 3’ or 5’ splice sites being used and their combination (**Figure 1A**). In the case of cassette exons, a complete alternative exon is included or skipped from the mature RNA, whereas in intron retention, an intron can be retained or removed. Alternative 3’ or 5’ splice sites result in a lengthening or shortening on either side of an exon, while alternative first and last exons rely on the usage of alternative transcription start sites and polyadenylation sites, respectively. Finally, mutually exclusive refers to two or more cassette exons that are used in exchange. In more complex transcriptomes such as in human, AS events are often not binary but occur in complex combinations, resulting in more than two possible splicing outcomes (Park et al. 2018).

**Figure 1.**
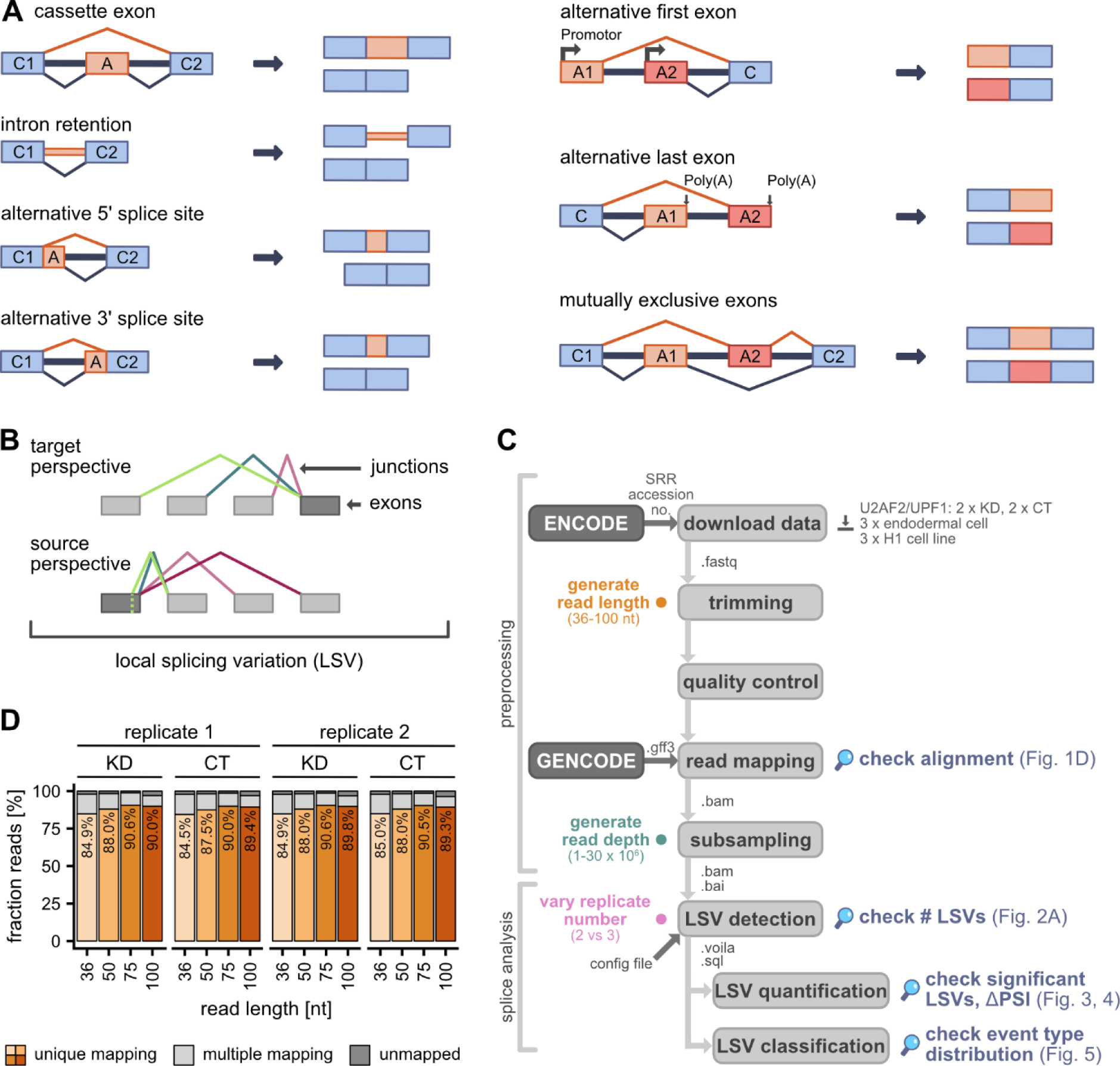
Exemplary alternative splicing (AS) analysis with MAJIQ. (A) Different types of AS events. Schematic displays pre-mRNA (left) and splice product(s) (right). pre-mRNAs include exons (rectangles), introns (thick lines) and splice junctions (thin lines). Blue elements always remain in the mature mRNA (constitutive, C), while orange and red elements are alternative (A). **(B)** Concept of local splice variation (LSV). An LSV describes a reference exon (dark grey) in the splice graph at which splice junctions (coloured) can start (source exon) or end (target exon). Exons are displayed as rectangles. **(C)** Workflow of pre-processing and AS analysis. First, the raw sequencing data was downloaded from ENCODE (fastq-dump), trimmed (Trimmomatic), checked for quality (FastQC) and then mapped to the human genome (STAR) with GENCODE gene annotation. Subsampling the resulting bam files simulated different sequencing depth (samtools view). AS analysis was performed using MAJIQ. Workflow includes work steps (light grey boxes), databases (dark grey boxes) and file formats (next to arrows). Coloured bullets indicate *in silico* variation in read length (orange), sequencing depth (sea green) and replicate number (pink). Magnifiers mark stages where influence of RNA-Seq characteristics was tested. **(D)** Longer reads show slightly better mappability. Proportion of uniquely mapped, multi-mapped and unmapped reads for four different read lengths (36–100 nt, light to dark orange) of the KD and control replicates of the *U2AF2* KD experiment.

Short-read RNA sequencing (RNA-Seq) is a popular experiment to elucidate AS in a transcriptome-wide manner. Depending on the sequencing strategy employed, read lengths vary between 25 and 250 nucleotides (nt), with sample sizes ranging up to 200 million (M) reads. As of February 2024, the database of the Encyclopedia of DNA Elements (ENCODE) Consortium offers more than 1,700 short-read RNA-Seq samples from human cell lines (Luo et al. 2020). A series of computational tools are available for AS analysis (Jiang and Chen 2021). The detection of AS events usually relies on junction-spanning reads, i.e., RNA-seq reads that span across the exon– exon junctions and thereby directly inform on the splicing status of the sequenced RNA molecule. As an example, the widely used algorithm Modeling Alternative Junction Inclusion Quantification (MAJIQ) (Vaquero-Garcia et al. 2016; Vaquero-Garcia et al. 2023) uses a splice site-centric approach based on local splicing variations (LSVs). In an LSV, all exon–exon junctions starting or ending in the same 5’ or 3’ splice site, respectively, are conjointly quantified (**Figure 1B**). This is particularly relevant for complex AS patterns with multiple overlapping AS events, reflected in a high ranking in previous benchmarking studies (Britto-Borges et al. 2021; Mehmood et al. 2020). Irrespective of the tool chosen, its performance is influenced by the characteristics of the RNA-Seq data. For instance, in gene expression analyses, more replicates generally increase statistical power (Schurch et al. 2016). The question of how to best set up an experiment for the detection of AS changes should therefore be considered carefully.

Here, we showcase the impact of different short-read RNA-Seq data characteristics on exemplary AS analyses using MAJIQ. We employ *in silico* variations of three public RNA-Seq datasets to test how different read lengths, read depths and replicate numbers influence the detection of significantly changing AS events (**Figure 1C**). The analysed datasets generated by the ENCODE Consortium (Luo et al. 2020) include the knockdown (KD) of the RNA-binding proteins U2AF2 and UPF1 as well as the differentiation of the human embryonic stem cell line H1 into endodermal cells, known to trigger widespread AS changes in the transcriptome (Chen et al. 2015; Kim and Maquat 2019; Zarnack et al. 2013).

## Results

### Read length marginally affects alignment efficiency

To test different RNA-Seq characteristics, we performed *in silico* variation on three standard short-read RNA-Seq datasets from the ENCODE Consortium (Luo et al. 2020). For read length and read depth, we used two experiments for *U2AF2* and *UPF1* KD in the immortalised human leukaemia cell line K562. Each comprised two replicates each for the KD and control condition, with on average 25.6 M (*U2AF2*) and 40.2 M (*UPF1*) paired-end (PE) reads per replicate, respectively. First, the raw reads were trimmed to four different read lengths (36–100 nt) and then aligned to the human reference genome using the splice-aware alignment algorithm STAR (Dobin et al. 2013) (**Figure 1C**). We found that 100-nt and 75-nt reads showed an almost identical fraction of uniquely mapping reads (89.6% and 90.4%, respectively), whereas shortening the reads to 36 nt reduced their mappability to 84.8% (mean over all replicates from *U2AF2* KD experiment; **Figure 1D**). Although all read lengths consistently achieved ≥ 80% uniquely mapped reads, the fraction of junction-spanning reads – which directly inform on splicing – went down for shorter reads (**Table 1**). Next, we generated eight different read depths by randomly subsampling the mapped 100- nt reads (1–30 million [M] PE reads per replicate; **Figure 1C**). Finally, to test the effect of replicates, we included an RNA-Seq dataset from the differentiation of the human stem cell line H1 into endodermal cells, consisting of three replicates for each condition with on average 51.7 M 100-nt paired-end reads. Here, we tested all possible combinations of two replicates and the three replicates. Altogether, the *in silico* variation yielded a total of 16 dataset variants, including four read length, eight read depth and four replicate number variants, for the subsequent AS analysis.

**Table 1.** Mean fraction and mean absolute number of junction-spanning reads among uniquely mapping reads over all replicates from *U2AF2* KD experiment.

| Read length | Percentage and absolute number of junction-spanning reads (mean of two replicates) |
| --- | --- |
| 36-nt | 20.9 (3,230,342) |
| 50-nt | 29.6 (4,753,796) |
| 75-nt | 46.1 (7,601,543) |
| 100-nt | 62.3 (10,182,618) |

### Read length and depth are critical to detect LSVs for subsequent quantification

To analyse AS changes, we employed the widely used algorithm MAJIQ and the associated visualisation tool VOILA (Vaquero-Garcia et al. 2016). In brief, the MAJIQ Builder first generates a unified graph representation of splice junctions present in the data (splicegraph) and defines all LSVs, i.e., 5’ or 3’ splice sites that engage in two or more alternative splice junctions and/or intron retention (LSV detection, **Figure 1C**). Next, the MAJIQ Quantifier calculates the relative abundance of each junction per LSV, i.e., the percent usage of the junction compared to all other junctions in the same LSV, termed percent selected index (PSI). To compare AS between conditions, the difference in PSI values for each junction (ΔPSI) is calculated together with the corresponding probability from a Bayes statistic (LSV quantification). Lastly, the VOILA Modulizer (Vaquero-Garcia et al. 2023) dissects the complex LSVs into distinct AS events that are contained therein (LSV classification). Starting with the *U2AF2* KD experiment, we performed individual MAJIQ analyses on the 16 dataset variants and compared the outcome at the three stages.

As a first metric, we assessed differences in LSV detection. MAJIQ uses a user-provided isoform annotation, but also detects LSVs *de novo* from the RNA-Seq reads. In both cases, LSVs are defined as present based on the junction-spanning reads in the data. We found that read length had a considerable impact on LSV detection (**Figure 2A**, left). Increasing from 36-nt to 100-nt reads more than doubled the number of LSVs (26,332 additional LSVs, 2.4-fold increase), as longer reads are more likely to span exon–exon junctions. LSV detection steadily improved with 50-nt and 75-nt reads and then stagnated with 100-nt reads (5,064 LSVs compared to 75-nt reads, 1.1-fold increase), indicating that saturation may be reached with reads over 100 nt.

**Figure 2.**
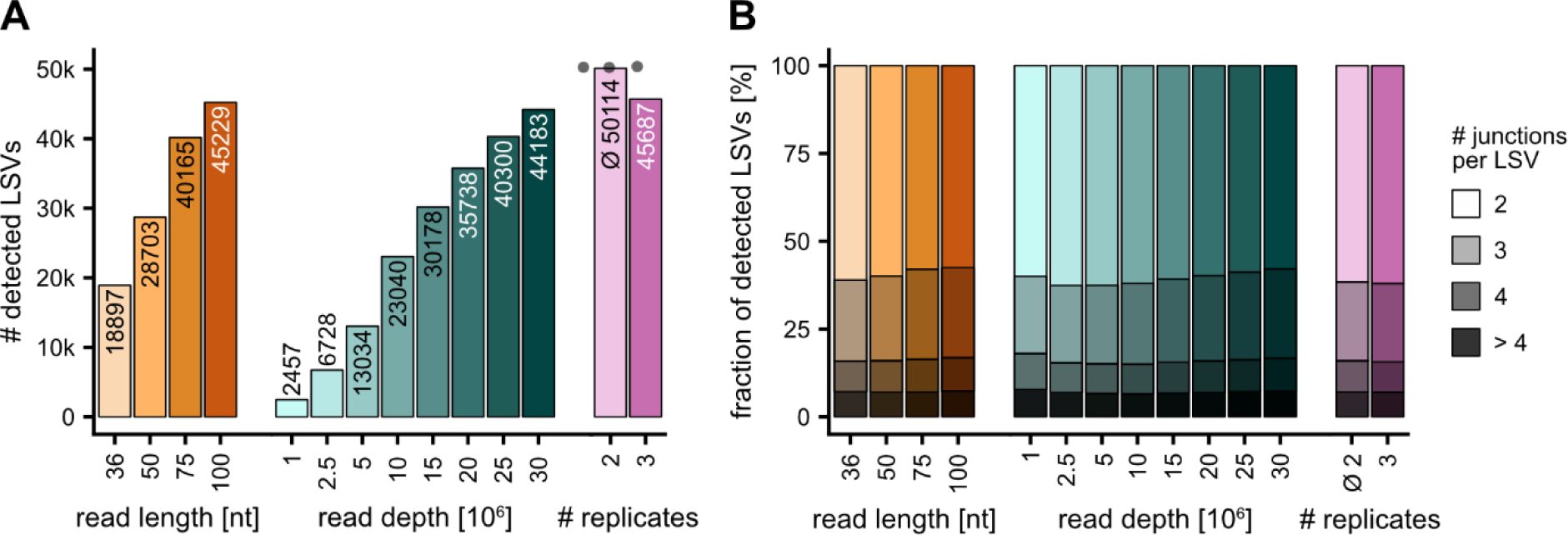
Longer reads and a higher read depth allow for better AS detection. **(A)** Number of LSVs detected by MAJIQ for different read lengths (36–100 nt, light to dark orange), read depths (1–30 M reads, light to dark sea green) and sample sizes (2 vs. 3 samples, light and dark pink). Individual datapoints and mean are shown for three possible combinations of two replicates. **(B)** LSV complexity. Fraction of LSVs with a given number of junctions therein for different read lengths (36–100 nt, light to dark orange), read depths (1–30 M reads, light to dark sea green) and replicate numbers (2 vs. 3 replicates, light and dark pink). Means of three possible combinations of two replicates are shown for all categories.

Varying the read depth displayed a comparable trend, with 30 M reads leading to 1.9-fold more detected LSVs than 1 M reads (**Figure 2A**, middle). Here, no onset of saturation was apparent yet even with the largest samples. Moreover, increasing read length or depth led to a progressive rise in the complexity of LSVs, reflected by the number of alternative junctions contained therein (**Figure 2B**). Of note, varying the number of replicates showed the opposite effect, as three replicates per condition yielded 4,427 (8.8%) fewer LSVs than just two replicates (**Figure 2A**, right). A reason might be that more replicates introduce a higher variability and therefore fewer LSVs are unambiguously identifiable. Repeating the analyses for the *UPF1* KD experiment yielded very similar results, with 2-fold more detected LSVs with increasing read length (from 35-nt to 100-nt; **Supplementary** Figure 1).

Overall, we concluded that the detection of LSVs strongly depends on the number and length of the RNA-Seq reads as longer reads or a higher read depth provide more information. Importantly, the LSVs that are detected in the first step set the boundary for the sensitivity of all following analyses as it restricts how many AS events can be identified as differentially regulated.

### Higher information content allows for more LSVs to reach significance

Next, we focused on LSVs that showed differential splicing between conditions. To define significant AS regulation, we asked for a minimum change in junction usage of 5% (ΔPSI ≥ 0.05) with a minimum probability of 90% (probability changing, P(|ΔPSI|>0.05) ≥ 0.9; **Figure 3A**). We found that longer reads allowed to identify more regulated LSVs and that these generally overlapped, indicating a higher sensitivity with increasing read length (**Figure 3B, C**). Again, this was corroborated in the *UPF1* KD experiment (**Supplementary** Figure 2). Interestingly, we observed an improvement not only in absolute numbers, but also at relative levels compared to the total LSVs detected. For instance, whereas 50-nt reads resulted in 952 regulated LSVs, corresponding to 3.3% of the LSVs detected in this dataset, this value increased to 2,671, equalling 5.9% of LSVs detected, for 100-nt reads. The same occurred with higher read depth (**Figure 3D, E**). While datasets with up to 5 M reads yielded hardly any regulated LSVs, the fraction of regulated LSVs steadily rose from 3.1% to 5.8% when moving from 10 M to 30 M reads. Together, these results indicated that even though many LSVs were already detected in the weaker datasets, they did not achieve sufficient coverage to reach significance. As expected, including more replicates led to more regulated LSVs, both in absolute and relative terms (**Figure 3F, G**). Altogether, these findings demonstrated that the sensitivity to identify regulated LSVs rises with all three RNA-Seq characteristics tested.

**Figure 3.**
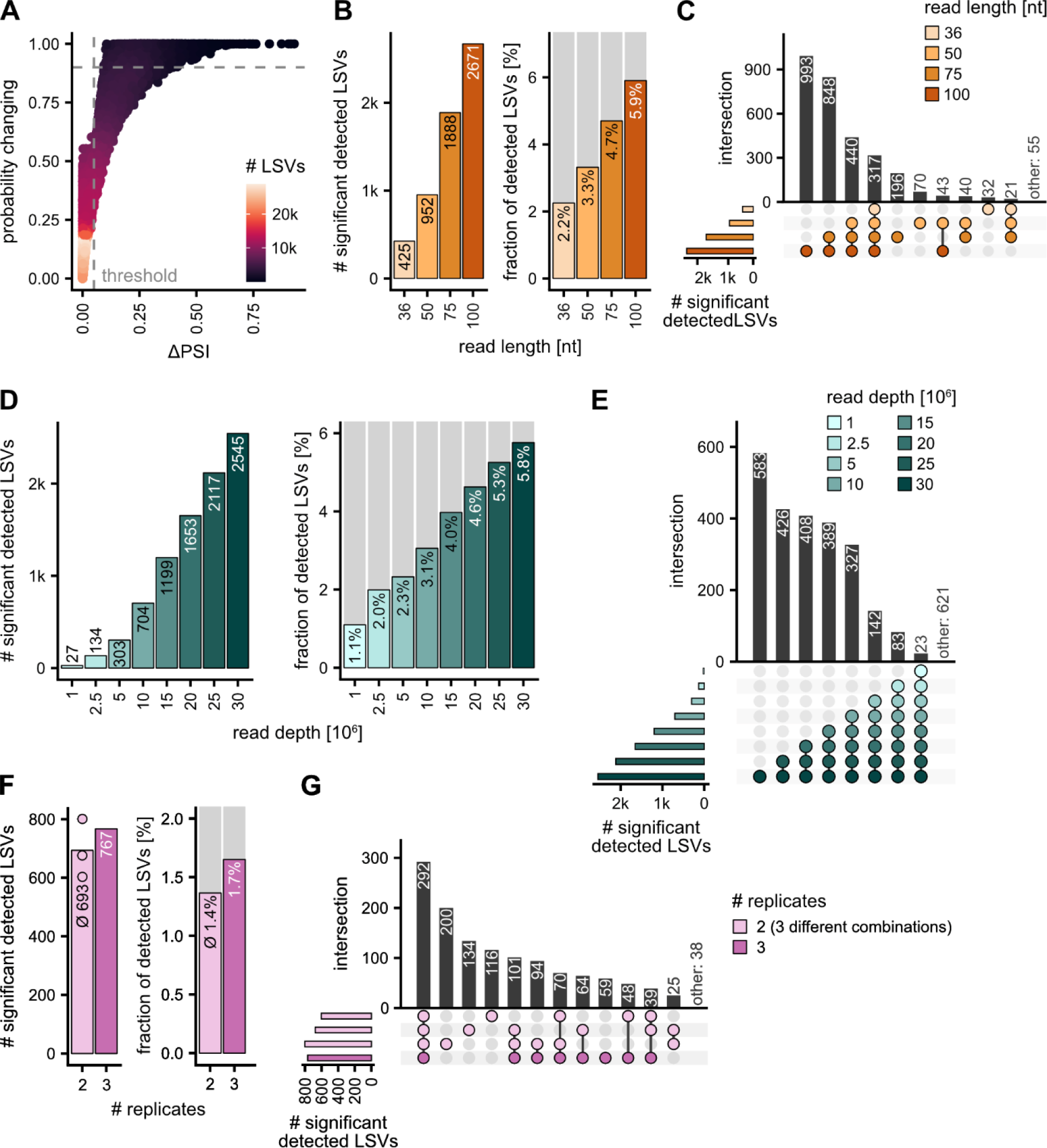
Higher information content leads to a greater sensitivity in the detection of significant differential splicing. (A) Maximum ΔPSI values are plotted against maximum probability changing (P(|ΔPSI| > 0.05)) for each LSV in the *U2AF2* KD dataset. Dashed grey lines show thresholds for an LSV to be considered significantly regulated (ΔPSI ≥ 0.05 and P(|ΔPSI| > 0.05) ≥ 0.9). Colour indicates density of LSVs (see legend). **(B–F)** Impact of read length (36–100 nt, light to dark orange), read depth (1–30 M reads, light to dark sea green) and replicate number (2 vs. 3 replicates, light and dark pink) on the identification of regulated LSVs. (B, D, F) Absolute number (left) and fraction (right) of significantly regulated LSVs retrieved for different read lengths (B), read depths (D), and replicate numbers (individual datapoints and mean for three combinations of two replicates are shown) (F). (C, E, G) UpSet plot shows overlap of regulated LSVs between test datasets. For replicate number, three possible combinations of two replicates are shown (light pink).

### MAJIQ returns conservative estimates for AS changes in scarce data

We next examined how the RNA-Seq library characteristics influenced the quantitative estimates of splicing changes between conditions. Given that an LSV consists of multiple junctions, we considered the maximum ΔPSI for each LSV. A comparison of absolute changes (|ΔPSI|) highlighted a trend whereby longer reads and higher read depths resulted in slightly larger absolute ΔPSI values (**Figure 4A, B, Supplementary** Figure 3A). The number of replicates also showed a small influence, with a slight tendency toward lower |ΔPSI| values for more replicates (**Figure 4C**). Importantly, a direct comparison of LSVs shared between datasets showed that the reduction in quantitative measurements mainly affected nonregulated LSVs, while regulated LSVs stayed close to the diagonal (**Figure 4D**, cyan vs. magenta linear regression line). This was consistently observed for both experiments and different read lengths, read depths and replicate numbers (**Figure 4D–F, Supplementary** Figure 3B). Moreover, the ΔPSI values were highly correlated for both nonregulated and regulated LSVs, indicating that the global reduction maintained the relative order between events. Thus, MAJIQ tends towards more conservative estimates of splicing changes and probabilities if information content in the data is low but maintains high precision for the regulated LSVs identified.

**Figure 4.**
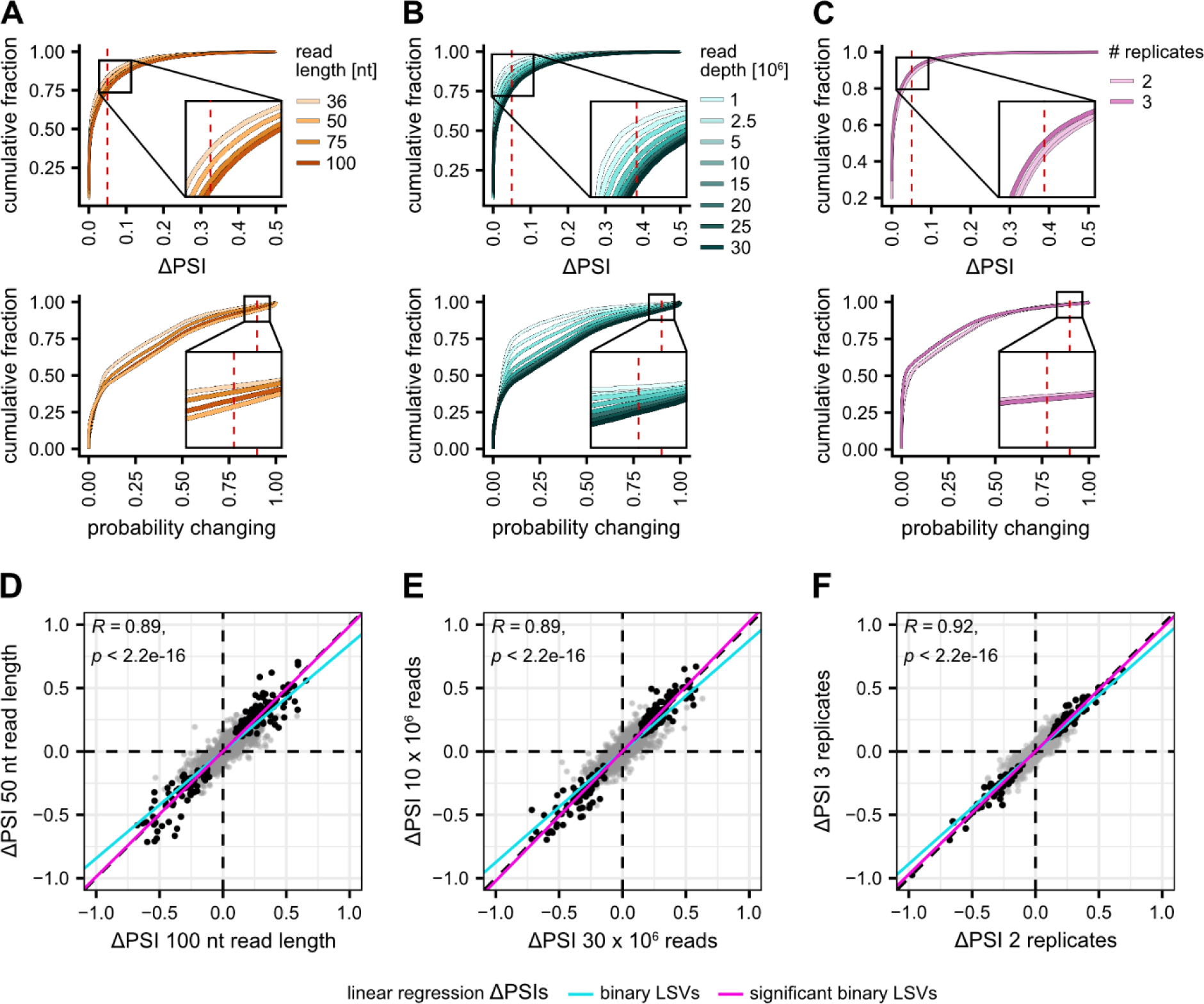
Quantitative estimates are downgraded for non-regulated events. (A–C) Shorter or less reads lead to smaller ΔPSI estimates. Cumulative fraction of ΔPSI and probability changing values for different read lengths (36–100 nt, light to dark orange), read depths (1–30 M reads, light to dark sea green), and replicate numbers (2 vs. 3 replicates, light and dark pink). For each LSV, the maximum ΔPSI and probability changing were considered. **(D–F)** Estimated changes for regulated LSVs are well reproducible. Exemplary comparison of ΔPSI values for shared LSVs between the 100-nt and 50-nt read datasets (D), the 30 M and 10 M read datasets (E), and using two and three replicates (F). LSVs that are significant in both datasets are shown in black. Magenta and cyan lines represent linear regression of ΔPSI values for regulated and all LSVs, respectively. Only binary LSVs are shown.

### Higher information content is required for AS detection in lowly expressed genes

To assess how the likelihood of detecting an AS change is linked to the expression of the underlying gene, we assessed the expression levels (transcripts per million, TPM) of genes harbouring regulated LSVs in the different dataset variants. Of note, regulated LSVs in highly expressed genes (TPM > 100) were efficiently detected with just 1 M reads per sample, whereas hardly any regulation was picked up in lowly expressed genes (TPM ≤ 25, **Figure 5A–D**). For example, at least 10 M reads were required to identify regulated LSVs in genes with TPM ≤ 5. These results can help to estimate the required read depth when interested in certain target RNAs. The number of replicates had no large effect on the identification of regulated LSVs in lowly expressed genes, but slightly fewer regulated LSVs were identified for the genes with very low TPM values (TPM ≤ 5) in the dataset with 3 replicates (**Supplementary** Figure 4).

**Figure 5.**
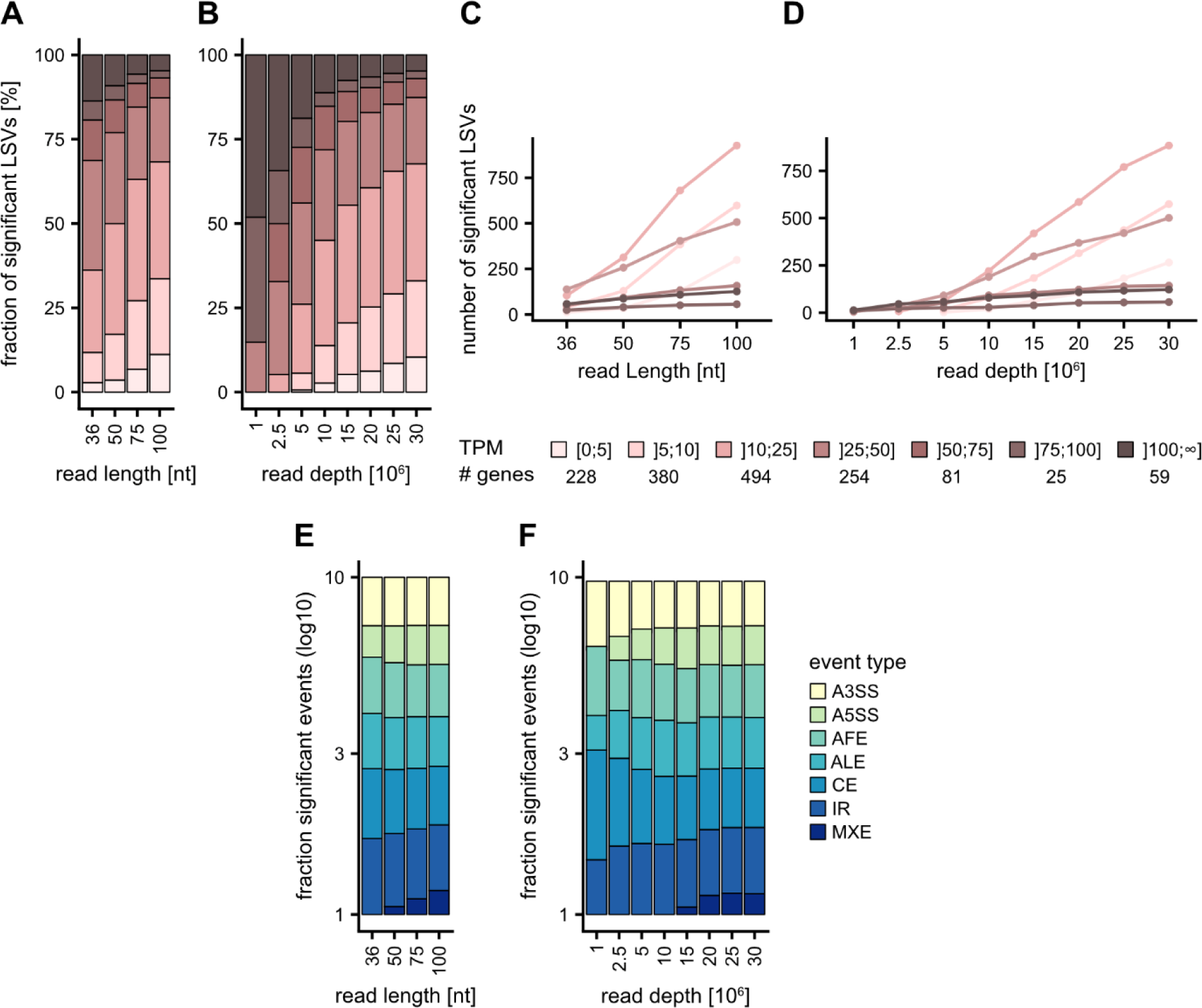
Detection of LSVs in lowly expressed genes requires data with higher information content. AS events involving three or more junctions benefit most from more complex datasets. (A–D) Fraction (A, B) and absolute number (C, D) of significant LSVs falling in bins with different TPM values for different read lengths (36– 100 nt) (A, C) and read depths (1–30 M reads) (B, D), ranging from light pink for low to brown for high TPM values. The number of genes in each bin is indicated below legend. **(E, F)** Fraction of regulated AS event types: alternative 3’ splice site (A3SS, yellow), alternative 5’ splice site (A5SS, light green), alternative first exon (AFE, green), alternative last exon (ALE, sea green), cassette exon (CE, petrol), intron retention (IR, blue), mutually exclusive exon (MXE, dark blue) for different (E) read lengths (36–100 nt) and (F) read depths (1–30 M reads).

Building on our previous findings, we further investigated the impact of the RNA-seq characteristics on the different types of AS events. The VOILA Modulizer allows to dissect LSVs into one or more AS events that are contained therein, such as cassette exons (CE), alternative splice sites (A5SS, A3SS) or intron retention (IR, **Figure 1A**). When considering regulated AS events, the detection of mutually exclusive exons (MXE) was most impaired by reduced read length, read depth or a limited number of replicates, although MXE were generally rare (**Figure 5E, F**). In addition, regulated A5SS events benefited from a higher information content in the data, which was achieved by increased read length and read depth.

## Discussion

Understanding the impact of RNA-Seq library characteristics on AS analysis is crucial to ensure the accuracy and reliability of gene expression studies. Here, we used *in sillico* variations of publicly available short-read RNA-Seq data to showcase the impact of read length, read depth, and replicates on differential splicing analysis. To this end, we present an exemplary analysis of standard RNA-Seq data as they are available in the widely used ENCODE database. Our results suggest that improving read length and depth will greatly benefit the detection of significantly regulated AS events.

In the context of differential gene expression, this has already been evaluated in several studies with varying resulting, ranging from one study concluding that just 200,000 reads are sufficient, but considering 9 samples (Baccarella et al. 2018), while another argues that most current experiments are undersequenced and recommends at least 60 M reads (Bass et al. 2019). As splicing detection relies particularly on junction-spanning reads, we expected the requirements for the RNA-Seq library characteristics to be higher for AS analysis compared to differential gene expression. Indeed, while it is suggested that 50-nt reads are sufficient for differential gene expression (https://knowledge.illumina.com/library-preparation/rna-library-prep/library-preparation-rna-library-prep-reference_material-list/000001243), we show here that reads of 100 nt or more are recommended for AS analysis. This is consistent with a previous study showing that 100-nt paired-end reads substantially increase splice site detection (Chhangawala et al. 2015).

The current ENCODE Guidelines and Best Practices for RNA-Seq experiments advise contributors to include at least two biological replicates of 30 M mapped reads or read pairs each (https://www.encodeproject.org/about/experiment-guidelines/). As in our analyses, neither the overall LSV detection nor the identification of regulated AS events saturated at this read depth, we recommend that this can be significantly expanded to realise the full potential.

Finally, in terms of replicates, we found only marginal differences in the detection of regulated AS events. We note that this is consistent with a previous study comparing sample sizes from 3 to 50 samples for different splicing analysis tools (Mehmood et al. 2020). Indeed, MAJIQ gave very similar results across all sample sizes, whereas most other tools improved substantially when more replicates were included. A previous study on differential gene expression found that six or more biological replicates should be used to achieve sufficient sensitivity and adequately control the false discovery rate (Schurch et al. 2016). This applied particularly to genes with reproducible but small changes in gene expression. Similarly, we found that increasing the information content of the data – through any of the factors tested – lowered the detection limit and improved the identification of AS events in lowly expressed genes. This is particularly relevant when studying regulatory genes with tightly controlled expression, such as transcription factors, which are often targeted by AS changes but are expressed at TPM < 20 in most tissues (Lambert et al. 2018; Soto et al. 2022).

### Recommendation

In conclusion, by evaluating factors such as read length, read depth, and replicates, researchers can optimise their experimental design. For a standard AS analysis, we recommend using RNA-Seq reads of 100 nt or longer and a read depth of at least 50 M reads per replicate. We note that paired-end information is not explicitly taken into account by MAJIQ but may be useful for other tools such as MISO (Katz et al. 2010). Three biological replicates per condition should be considered as an absolute minimum to allow for sufficient statistical power. We recommend starting with more replicates, as it is common for one or more replicates to be removed in later analyses due to quality issues. Overall, these settings will facilitate read alignment, a high number of splicing events detected and a good sensitivity for identifying regulated alternative splicing.

## Materials and Methods

### Experimental data

All analysed RNA-seq datasets were taken from the ENCODE consortium (Luo et al. 2020). Two experiments were for *U2AF2* or *UPF1* knockdown (KD) in the immortalised human leukaemia cell line K562 (ENCODE IDs: *U2AF2*, ENCSR904CJQ; *UPF1*, ENCSR251ABP). In brief, the expression of *U2AF2* and *UPF1* was depleted using small hairpin RNAs (shRNAs). Non-targeting shRNAs served as controls (CT). The experiments had two biological replicates for each condition. To compare the influence of sample size, we further included two total RNA-Seq datasets for the human stem cell line H1 (ENCSR588EJX) and human differentiated endodermal cells (ENCSR266XAJ), each with three biological replicates. These data have also been used as short-read RNA-Seq reference in a community challenge of the GENCODE Long-read RNA-seq Genome Annotation Assessment Project (LRGASP) Consortium in 2021 (Pardo-Palacios et al. 2023). All samples were generated by Illumina short-read sequencing with 100-nt paired-end reads and contained on average 41.0 million read pairs per replicate.

### Pre-processing and *in silico* variation of sequencing data

All RNA-Seq datasets were downloaded in FASTQ format from the SRA database using fastq-dump (sra-toolkit version 3.0.0; https://github.com/ncbi/sra-tools/wiki).

Trimmomatic (version 0.39) (Bolger et al. 2014) was used to remove adapter sequences from the 3’ ends with `ILLUMINACLIP:TrueSeq3- PE.fa:2:30:10:8:true` (*U2AF2* and *UPF1* KD) and `ILLUMINACLIP:NexteraPE-PE.fa:2:30:10:8:true` (H1 differentiation). Sliding window quality trimming was performed with `--SLIDINGWINDOW:4:20`.

Reads with a length < 100 nucleotides (nt) were sorted out (`--MINLEN:100`). To obtain different read lengths, all reads were then cropped to 100, 75, 50 and 36 nt (`--CROP:desiredLength`).

All FASTQ files were subjected to quality control using FastQC (version 0.11.9; https://www.bioinformatics.babraham.ac.uk/projects/fastqc/). Then, the reads were aligned to the human reference genome (version GRCh38.p13) with GENCODE gene annotation (version 41) (Frankish et al. 2019) using STAR (version 2.7.10a) (Dobin et al. 2013). For mapping, the paired-end option (as collection) was used and `-- sjdbOverhang` was set to read length - 1. Only uniquely mapping reads (`-- outFilterMultimapNmax 1`) with up to 4% of mismatches per mapped read length (`--outFilterMismatchNmax 999 --outFilterMismatchNoverLmax 0.04`) were processed further. Unaligned reads were excluded from the BAM output.

To simulate different sequencing depths, the mapped reads of the *U2AF2* KD dataset were subsampled to 30 M, 25 M, 20 M, 15 M, 10 M, 5 M, 2.5 M, and 1 M reads using `samtools view` (version 1.6) (Li et al. 2009).

### Alternative splicing analysis

AS analysis was performed using MAJIQ (version 2.3) (Vaquero-Garcia et al. 2016; Vaquero-Garcia et al. 2023). First, the MAJIQ Builder was used for LSV detection (`majiq build`). It was provided with the GENCODE gene annotation (version 41) in GFF3 format and a configuration file containing the BAM file paths, information on read length, strandedness (“reverse”) and the reference genome. The experiments were grouped based on KD compared to CT and the stem cell line H1 compared to differentiated endodermal cells. To generate different sample sizes, the MAJIQ Builder was run multiple times with configuration files that had different numbers of replicates specified.

Next, the MAJIQ Quantifier (`majiq deltapsi`) was run for quantifying differences in junction usage (delta percent selected index, ΔPSI) on the MAJIQ files obtained from the MAJIQ Builder. To visualize the outcome, the SQL file from the MAJIQ Builder, containing the splicegraph, and the VOILA files from the MAJIQ Quantifier were used as input for VOILA. `voila tsv` was executed with the following parameter settings:

--show-all --threshold 0.05 --changing-between-group-dpsi 0.05

--non-changing-between-group-dpsi 0.05.

For AS event classification, the VOILA Modulizer (voila modulize, also called voila categorize in newer versions) was run with the following parameter settings: --changing-between-group-dpsi 0.05 --non-changing-between-group-dpsi 0.05 --changing-between-group-dpsi-secondary 0.025 --show-all.

The output from `voila tsv` and `voila modulize` was then further analysed in R, where we filtered for significantly regulated LSVs (ΔPSI ≥ 0.05 and P(|ΔPSI| > 0.05) ≥ 0.9 probability changing, **Figure 3, 5**). Since an LSV can have multiple junctions with different ΔPSI and probability changing values, we chose the maximum of these values for each LSV.

Regulated AS events (**Figure 5**) with two splice junctions (A3SS, A5SS, AFE, ALE, IR) were considered regulated if at least one junction met the regulation criteria. For cassette exons (CE) or mutually exclusive exons (MXE) with four splice junctions, we considered them as regulated if either both junctions originating from C1 or both junctions originating from C2 met the thresholds of ΔPSI ≥ 0.05 and P(|ΔPSI| > 0.05) ≥ 0.9 (**Figure 1A**). The less common splicing events described – MAJIQ classifies more than 10 different types – were left out for simplicity.

### Calculation of TPM values

HTSeq (`htseq-count`; version 2.0.2) (Anders et al. 2015) was run to count the number of reads mapping to specific exons for the *U2AF2* KD experiment (100 nt long reads), for the human stem cell line H1 and for the human differentiated endodermal cell dataset. BAM files and gene annotation were passed with the following options: `--stranded reverse -–type exon -–nonunique all`. The resulting counts were averaged across all replicates in order to compute Transcripts Per Million (TPM) values. First, mean counts are divided by gene lengths to obtain Reads Per Kilobase (RPK) values. Subsequently, the RPK value was divided by the sum of all mean counts. As gene length the total length of all annotated merged exons was used (GenomicFeatures, version 1.54.4, `exonsBy`, `by = “gene”`; GENCODE gene annotation version 41; GenomicRanges, version 1.54.1, `reduce`). Only TPM values of genes with regulated LSVs were considered for analysis.

### Data availability

The analysed RNA-seq data are available in the ENCODE database (https://www.encodeproject.org/) with the following accession numbers: *U2AF2* KD in human K562 (ENCODE ID: ENCSR904CJQ), *UPF1* KD in human K562 (ENCODE ID: ENCSR251ABP), human stem cell line H1 (ENCODE ID: ENCSR588EJX), and human differentiated endodermal cells (ENCODE ID: ENCSR266XAJ).

The computational code for this study is available on GitHub at https://github.com/ZarnackGroup/Ladwig_et_al_2024.

## Acknowledgements

The authors would like to thank Yoseph Barash and all members of the Zarnack group for valuable discussions and Mario Keller for help with MAJIQ.

## Author contributions

A.L. analysed data. M.K., K.Z., designed studies and interpreted data. A.L., M.K., K.Z. wrote the manuscript and made the figures.

